# Pangenome-resolved polymorphic transposable elements reveal human regulatory evolution in action

**DOI:** 10.64898/2026.09.24.754167

**Authors:** Juan Jiang, Xiaoyu Zhuo, Ronghan Li, Zheng Dong, Wenjin Zhang, Daofeng Li, Juan F. Macias-Velasco, Human Pangenome Reference Consortium, Ting Wang

## Abstract

Transposable elements are a major source of gene-regulatory functions. Yet this contribution is understood largely from ancient insertions already fixed in the genome: the successes, seen long after the fact. The polymorphic phase before fixation or loss remains poorly resolved because such alleles could be scored only as present or absent. We resolved segregating transposable-element alleles as sequences, epigenetic states, and transcript components using 231 haplotype-resolved human pangenomes with matched long-read multi-omics. Their origins extend beyond recent transposition; their regulatory consequences depend on methylation and transcript context and are often missed by gene-level analyses; they provide mechanisms at trait-associated loci, while selection favors element presence and absence alike. Regulatory innovation before fixation is therefore not one-way accumulation, but a mixture of gain, remodeling, and loss.

## MAIN

Transposable elements (TEs) are major architects of genome evolution and regulatory innovation(*1–3*). By proliferating within the host genome, TEs generate lineage-specific sequence diversity and supply abundant genetic material that can be co-opted into host regulatory programs. TE-derived sequences can act as cis-regulatory elements (cCREs)(*4–8*), remodel transcript structures through alternative promoters, splice sites and polyadenylation signals(*9, 10*), and exert post-transcriptional control through RNA structure and RNA-binding protein interactions(*11–13*). However, our view of these processes remains largely retrospective. Most mechanistic examples involve ancient insertions that are already fixed in reference genomes and represent the products of evolutionary processing. How TE-derived regulatory functions arise, generate molecular and phenotypic variation, and become shaped by evolutionary processes while the underlying alleles remain polymorphic therefore is poorly understood.

Polymorphic TEs (poly-TEs) provide a natural system to address this question because TE-present and TE-absent allele states coexist within contemporary populations. Across humans and other species, previous studies have established poly-TEs as an important source of genetic variation by cataloging insertions across populations(*14–17*), linking subsets to variation in gene expression and splicing(*16, 18–22*), and to traits and disease(*16, 23–26*), and identifying individual TE alleles that contribute to local adaptation(*27–29*). However, most previous studies have relied on short-read sequencing and linear-reference representations, leaving many non-reference insertions, repetitive internal sequences and allele-specific molecular states unresolved. Consequently, poly-TEs have often been analyzed as insertion-presence markers rather than as sequence-resolved haplotypic alleles, limiting the ability to determine which alleles acquire regulatory activity and through which mechanisms that activity is realized. Long-read sequencing(*30, 31*), haplotype-resolved pangenomes(*32, 33*), native DNA methylation profiles(*34, 35*) and full-length transcriptomes now make it possible to resolve these alleles and their regulatory states.

Here, we address this question by integrating the Human Pangenome Reference Consortium’s (HPRC) Release 2 (HPRC2) haplotype-resolved pangenome panel with matched long-read methylation profiles, full-length transcriptomes(*36*), molecular quantitative trait loci (QTL) mapping, trait-associated loci and population genetics analyses (table S1). This framework places poly-TE alleles in their native sequence and haplotype contexts, allowing their molecular states and regulatory consequences to be interpreted together. We show that present-day poly-TEs arise through diverse evolutionary routes and generate regulatory variation across multiple DNA- and RNA-level mechanisms. By linking these alleles to molecular QTLs, trait-associated haplotypes and recent selection, our study reveals how TE-derived regulatory variation is generated, realized and evolutionarily processed before fixation or loss, providing a population-level view of regulatory innovation in action.

## Results

### Human polymorphic TE variation extends beyond recent transposition

Using the decomposed HPRC2 pangenome Minigraph-Cactus graph(*36*), which is represented in GRCh38 coordinates and incorporates CHM13 together with 231 diploid human assemblies, we first annotated 65,906 TE-containing structural variants (SVs) across the graph (table S2). Among these, 37,244 TE loci were absent from GRCh38 and 35,087 were absent from CHM13 but present in at least one HPRC2 haplotype, highlighting the additional TE diversity captured by the pangenome. Of the 65,906 loci, 64,783 segregated across the 231 diploid genomes, whereas the remaining 1,123 were invariant within the HPRC2 cohort. We refer to these segregating loci as polymorphic TEs (poly-TEs). Of these, 23,638 loci exhibited hallmarks of recent mobilization and were classified as mobile element insertions (MEIs), whereas 41,145 lacked canonical mobilization signatures and were classified as non-MEI poly-TEs (Fig. 1A). In parallel, 4,451,580 TE loci were fixed across all 231 diploid genomes. Comparison with MEIs identified from sample-matched (n = 223) high-coverage short-read sequencing in the 1000 Genomes Project (1000G) revealed that nearly half of the pangenome-resolved MEIs were absent from 1000G’s short-read call set (Fig. 1B). These HPRC2-specific MEIs were overrepresented by longer MEIs (L1, SVA, and HERV-K), and enriched in structurally complex regions and low-frequency alleles, indicating that pangenomes recover a large subset of MEIs inaccessible by short reads (Fig. 1C). Beyond canonical MEIs, the pangenome uncovered a substantial class of non-MEI poly-TEs. Non-MEI poly-TEs represent SVs containing TE fragments, most of which retained simple TE-rich architectures: 70.9% were composed predominantly of TE sequence (TE fraction >50%), and 58.5% harbored only a single TE. These non-MEI poly-TEs spanned a broad allele-frequency spectrum, showed intermediate sequence divergence between MEIs and fixed TEs, and were distributed across both young and old TE families (fig. S1A and B).

**Fig. 1.**
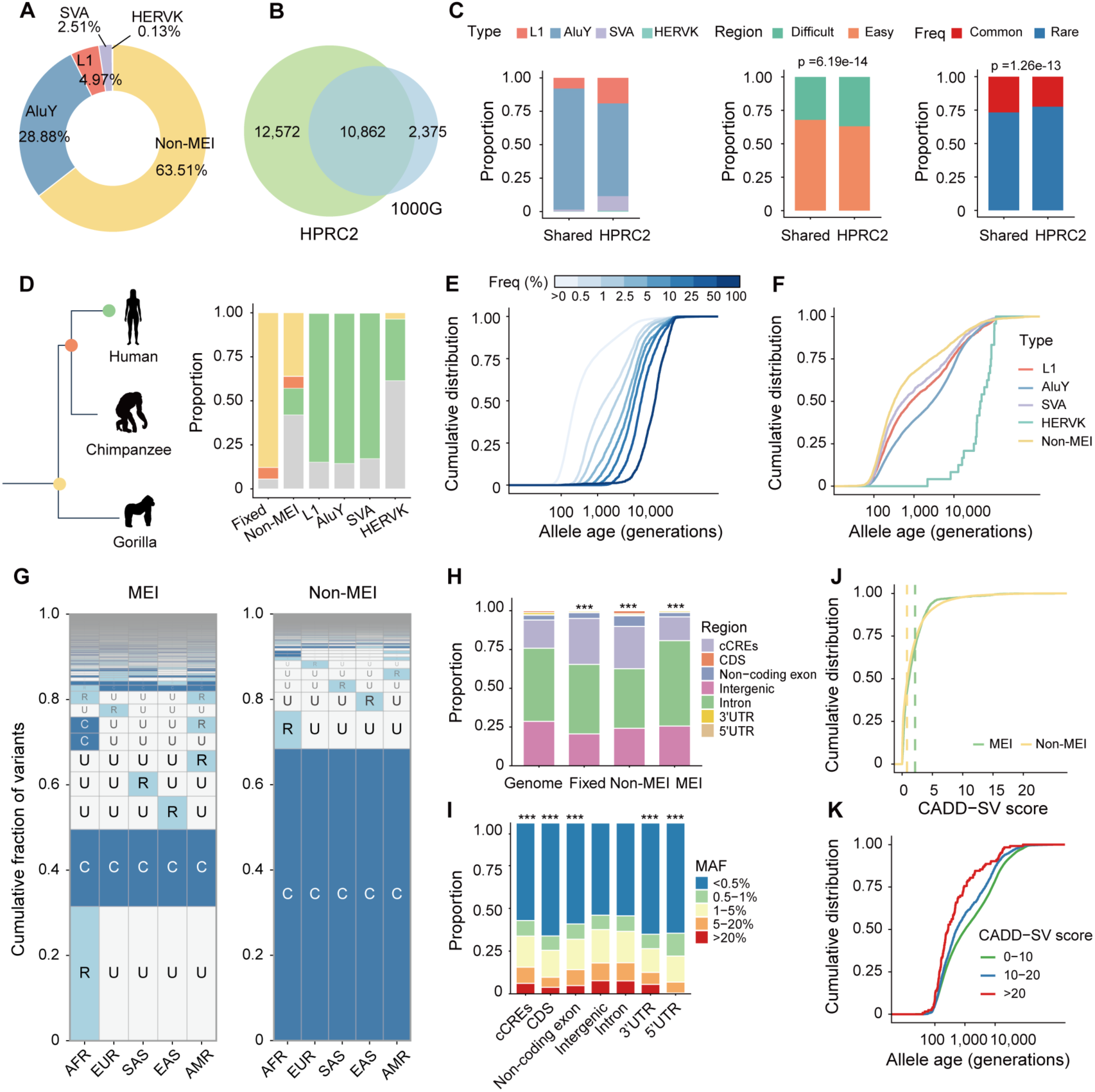
Pangenome resolution reveals diverse evolutionary origins of polymorphic TE alleles. (**A**) Proportion of different TE types among polymorphic TEs (poly-TEs). Non-MEI poly-TEs denote poly-TEs excluding mobile element insertions (MEIs; including L1, AluY, SVA and HERV-K insertions). (**B**) Overlap of MEIs identified in HPRC2 and sample-matched 1000 Genomes Project (1000G) data. (**C**) Comparison between HPRC2-only and HPRC2-1000G shared poly-TEs. Left, proportions of MEIs and non-MEI poly-TEs located in GIAB difficult and easy regions; middle, TE frequency distributions; right, TE type composition. Statistical significance was assessed using two-sided Fisher’s exact tests. (**D**) Phylogenetic states of different TE types. Species silhouettes were obtained from PhyloPic. Yellow, shared by human and gorilla; orange, shared by human and chimpanzee; green, human specific. (**E**) Allele age distribution of poly-TEs stratified by derived allele frequency. (**F**) Allele age distribution of human-specific poly-TEs. Ages in e and f are plotted on a log_10_ scale. (**G**) Geographic distribution of poly-TEs across 1000G superpopulations, plotted with geovar package(*95*). C, common TEs with TE frequency > 5%; R, rare TEs with 0 < TE frequency ≤ 5%; U, unobserved TEs with TE frequency = 0. AFR, African; EUR, European; SAS, South Asian; EAS, East Asian; AMR, Admixed American. (**H**) Genomic-region distribution of fixed TEs, MEIs and non-MEI poly-TEs compared with the genome background. Regions were assigned using a mutually exclusive priority order: CDS, 5′UTR, 3′UTR, non-coding exon, cis-regulatory elements (cCREs), intron and intergenic. For statistical testing, CDS, UTRs, non-coding exons and cCREs were grouped as functional regions and compared against intronic/intergenic regions. Each TE class was compared with the genome background using chi-square tests, and P values were adjusted using the Benjamini–Hochberg method. ***FDR < 0.001. (**I**) Minor allele frequency (MAF) of poly-TEs across functional genomic regions. Fisher’s exact tests were used to compare the proportion of rare alleles (MAF ≤ 5%) in each region against intergenic regions, with all remaining alleles grouped as MAF > 5%. P values were adjusted using the Benjamini–Hochberg method. (**J**) Distribution of Combined Annotation Dependent Depletion for Structural Variants (CADD-SV) scores(*96*) for poly-TEs. The dashed line denotes the median value. (**K**) Derived allele age distribution of poly-TEs stratified by CADD-SV score.

Outgroup polarization using chimpanzee and gorilla assigned ancestral states to nearly 80% of fixed TEs and MEIs, but only about half of non-MEI poly-TEs. Fixed TEs were predominantly shared with two outgroups, whereas MEIs were predominantly human-specific TE insertions (Fig. 1D). A small number of MEIs, including 8 L1, 67 AluY, 4 SVA and 3 HERV-K insertions, were also present in the chimpanzee genome, consistent with the persistence of ancestral MEI polymorphisms through incomplete lineage sorting, as previously described for PtERV1 elements retained in chimpanzee and gorilla but absent from humans(*37*). In contrast, non-MEI poly-TEs showed mixed origins, with approximately 74% shared with one or both outgroups and 26% being human specific, indicating that non-MEI poly-TEs comprise both ancestral segregating TE alleles and human-specific TE-associated SVs (Fig. 1D). Genealogy-based age estimates further placed poly-TE alleles on a within-human timescale, showing that allele age generally increased with frequency but remained highly variable among frequency-matched alleles (Fig. 1E). Although most non-MEI poly-TEs showed greater sequence divergence from consensus than MEIs (fig. S1B), human-specific non-MEI poly-TE alleles were younger than human-specific canonical MEIs (Fig. 1F), suggesting that TE-associated polymorphism is generated not only by de novo mobilization, but also by rapid structural remodeling of pre-existing TE-containing haplotypes.

The geographic distribution of TE-presence alleles further reflected evolutionary history and recent demography. Whereas most MEI families were enriched for geographically restricted alleles, HERV-K MEIs and non-MEI poly-TEs were more broadly shared across populations (Fig. 1G and fig. S1C). African individuals carried greater poly-TE allelic diversity and heterozygosity, while non-African individuals showed higher homozygosity, consistent with non-African bottlenecks and population-specific drift (fig. S1D). At the same time, the total TE allele counts combining heterozygous and homozygous insertions were comparable across all five superpopulations, suggesting that demographic history reshaped the distribution of TEs across genotype states rather than producing major differences in overall per-individual TE allele counts (fig. S1D). Archaic introgression further contributed 457 poly-TEs (table S2), adding a fourth source of poly-TEs in present-day humans, including Neanderthal- and Denisovan-derived alleles near notable loci such as *OCA2*, *STAT2*, *ABO*, *TSHR* and *PGR*.

Poly-TEs also bore signatures of functional constraint. They were predominantly intronic and intergenic and depleted from functional elements (exons or cCREs) (Fig. 1H). Those overlapping functional elements segregated at lower frequencies, consistent with purifying selection against disruptive TE variation (Fig. 1I). MEIs showed higher predicted deleteriousness than non-MEI poly-TEs (Fig. 1J), and higher deleterious scores were associated with younger allele ages (Fig. 1K), consistent with recent disruptive alleles having had less time to be removed by purifying selection. Together, these analyses show that human poly-TEs are not a single class of recent mobilization but represent a heterogeneous evolutionary spectrum arising from ancestral segregation, recent mobilization, structural remodeling of older repeats and archaic introgression, and subsequently shaped by demographic history and purifying selection.

### Polymorphic TEs show genomic-context-dependent DNA methylation and reshape local epigenetic landscapes

To understand how epigenetic state constrains or enables the regulatory potential of poly-TEs, we first characterized their DNA methylation landscape. Poly-TEs were generally more highly methylated than fixed TEs, with MEIs showing the strongest hypermethylation (fig. S2), consistent with robust host silencing of recently active insertions. However, MEI methylation was not uniform across element bodies (Fig. 2A). Alignment of methylation profiles to subfamily consensus sequences revealed family-specific patterns, including reduced methylation near the L1HS ORF1–ORF2 boundary, which encompasses the internal ribosome entry site; reduced methylation between the R and U5 regions of LTR5_Hs, near the retroviral transcription termination site; and focal hypomethylation near the 3′ ends of L1HS and SVA elements (Fig. 2A). The region encompassing the HERV-K (HML-2) transcription start site, located at the U3– R boundary within the LTR, showed the largest methylation difference between rare (allele frequency ≤ 0.05) and common (allele frequency > 0.05) LTR5_Hs insertions (Fig. 2A). Thus, MEI silencing is structured rather than uniform, leaving subregions with potential regulatory or transcriptional relevance.

**Fig. 2.**
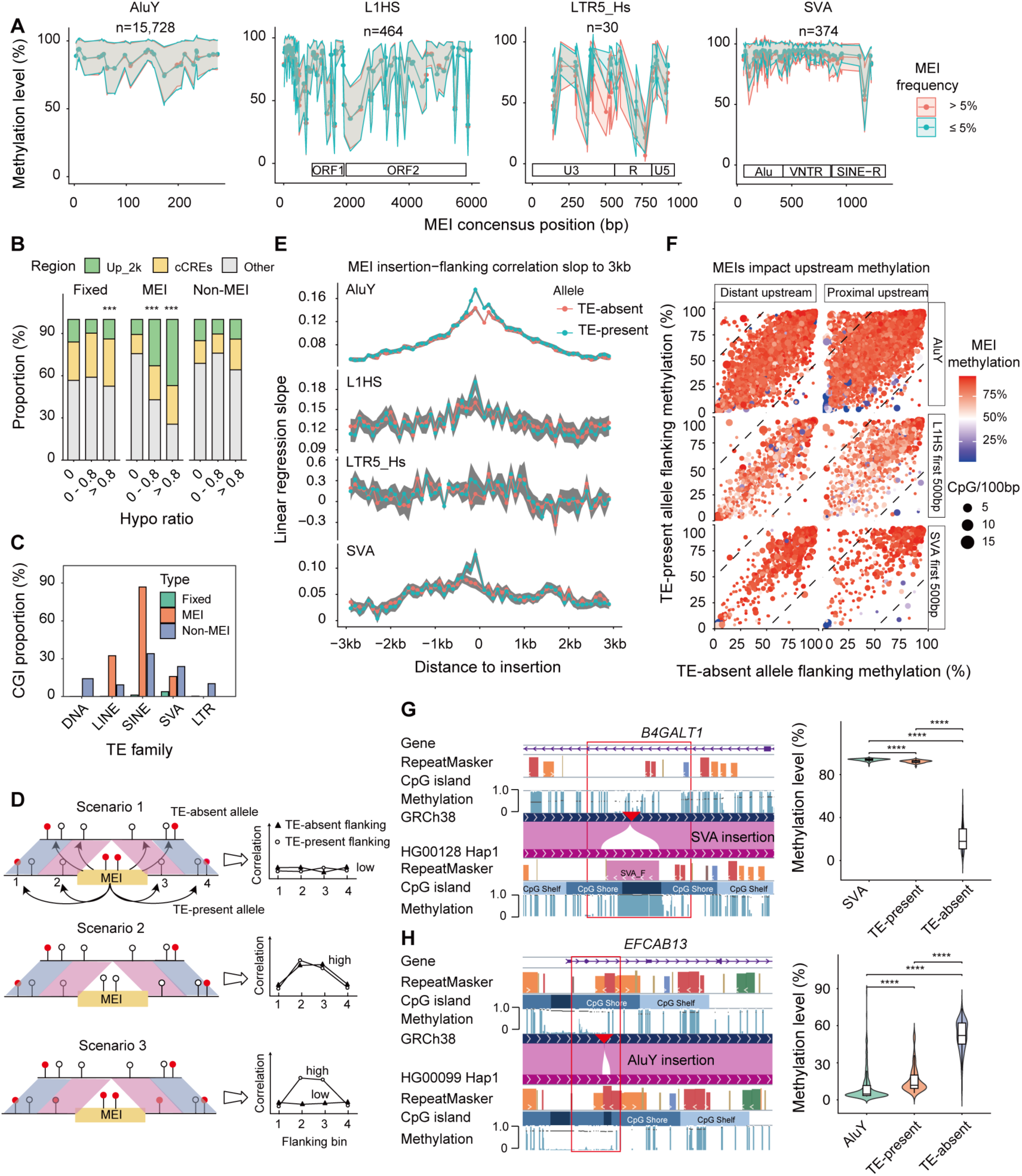
DNA methylation of polymorphic TEs and its effect on the surrounding epigenetic landscape. (**A**) MEI methylation profiles aligned to TE consensus sequences. Lines show per-position mean CpG methylation across MEI copies; shading areas indicate 95% standard error of the mean. (**B**) Enrichment of hypomethylated TEs in promoter-proximal regions and cis-regulatory elements (cCREs) within each poly-TE class. Hypo ratio = 0 indicates no hypomethylated (methylation level < 0.3) carriers; 0 < hypo ratio ≤ 0.8 includes TEs with at least one hypomethylated carrier haplotype, including single-carrier TEs, but for multi-carrier TEs the hypomethylated fraction did not exceed 0.8; and hypo ratio > 0.8 indicates hypomethylation in >80% of carrier haplotypes, calculated only for TEs carried by at least two haplotypes. Up_2k, 2 kb upstream of transcription start sites. Enrichment was tested relative to TEs without hypomethylated carriers (hypo = 0) within the same TE class using Fisher’s exact test. ***P < 0.001. (**C**) Fraction of TEs overlapping CpG islands (CGI) annotated in haplotype-resolved assemblies. (**D**) Schematic of three relationships between MEI and flanking methylation: scenario1, independent methylation, scenario2, shared local epigenetic context, and scenario3, insertion-associated remodeling of nearby methylation. (**E**) Association between MEI body methylation and flanking methylation across distance from the insertion. The y axis shows the linear regression slope between mean MEI methylation and methylation in each 100-bp flanking bin, calculated separately for TE-absent and TE-present haplotypes and shown by MEI family. Shading areas indicate 95% standard error of the mean. (**F**) Allele-specific flanking methylation at individual MEI loci. The x and y axes show flanking methylation on TE-absent and TE-present haplotypes, respectively; color indicates MEI methylation and point size indicates CpG density. Dashed lines mark >50% methylation differences between haplotypes. Distant upstream flanks (900 bp to 1 kb) and proximal upstream flanks (0–100 bp) are shown for AluY, proximal L1HS and proximal SVA. (**G**) and (**H**) Representative MEIs associated with increased (G, hypermethylated SVA) or reduced (H, hypomethylated AluY) methylation in the 300-bp flanking regions. Left, HPRC epigenome browser view of MEI and flanking methylation, with tracks for gene annotation, RepeatMasker, CpG islands, and PacBio methylation and sequence alignment,. Red box highlighted the MEI and associated methylation change; right, methylation comparison among TE body, TE-absent and TE-present haplotypes using two-sided Wilcoxon rank-sum tests. ****P < 0.0001.

Despite this pervasive silencing, a subset of poly-TEs escaped methylation repression. In total, 1,258 poly-TEs were hypomethylated (methylation level < 0.3) in at least one carrier. For some loci, this state recurred across individuals: 51 MEIs and 540 non-MEI poly-TEs were hypomethylated in more than 80% of carriers. Hypomethylated MEIs were strongly enriched in promoter-proximal regions and cCREs, with enrichment positively correlated with the percentage of hypomethylated haplotypes; more than 70% of consistently hypomethylated MEIs overlapped these annotations (Fig. 2B). This pattern may be particularly relevant for AluY insertions, approximately 90% of which form CpG island, significantly higher than fixed elements (Fig. 2C). Together, these findings identify a subset of young MEIs that repeatedly escapes host silencing in regulatory environments, allowing their CpG-rich sequences to remain hypomethylated and potentially acquire regulatory activity.

We next asked whether MEI methylation is shaped by the pre-existing local epigenetic environment and whether MEI insertion can, in turn, alter methylation in neighboring sequences. To distinguish these effects, we correlated MEI-body methylation with flanking methylation measured either on the TE-present haplotype or at the corresponding locus on the TE-absent haplotype. Methylation on the TE-absent haplotype served as a proxy for the local epigenetic state before insertion. Weak correlations with flanking methylation on both haplotypes indicate that MEI methylation is largely independent of local context (Fig. 2D, Scenario 1). Similar correlations on TE-present and TE-absent haplotypes indicate that MEI methylation primarily reflects the pre-existing local environment (Scenario 2), whereas stronger correlations on the TE-present haplotype indicate insertion-associated remodeling of nearby methylation (Scenario 3). AluY insertions showed significant correlations with flanking methylation on both haplotypes, indicating strong sensitivity to local epigenetic context. These correlations were nevertheless stronger on TE-present haplotypes, suggesting that AluY insertions can additionally reshape nearby methylation (Fig. 2E). By contrast, L1HS and SVA methylation showed little correlation with flanking methylation on TE-absent haplotypes but increased correlation with the upstream 100 bp on TE-present haplotypes, consistent with insertion-associated remodeling. LTR5_Hs methylation remained only weakly correlated with flanking methylation on either haplotype, suggesting relative independence from the surrounding methylation state (Fig. 2E).

We next examined whether MEI insertions were associated with directional changes in flanking methylation. For each locus, we plotted methylation on the TE-present haplotype against methylation at the corresponding region on the TE-absent haplotype, with MEI methylation indicated by color (Fig. 2F). Loci with similar methylation on the two haplotypes fall along the diagonal, whereas loci above or below the diagonal show increased or decreased methylation, respectively, on the TE-present haplotype. Allele-specific differences were minimal in the distal upstream region, represented by the 100-bp bin located 1 kb from the insertion. By contrast, the 100 bp immediately upstream of the insertion showed a pronounced increase in loci with methylation more than 50 percentage points higher on the TE-present haplotype. The number of such loci increased from 14 in the distal bin to 210 in the proximal bin for AluY, from 3 to 20 for L1HS and from 3 to 24 for SVA. Loci with methylation more than 50 percentage points lower on the TE-present haplotype were much less common and changed only modestly between distal and proximal regions, from 19 to 17 for AluY, from 3 to 6 for L1HS and from 2 to 4 for SVA. The increases in L1HS and SVA were largely driven by elements whose first 500 bp were hypomethylated, including four L1HS and three SVA insertions. These changes tracked MEI methylation state: hypermethylated insertions were associated with increased nearby methylation, whereas hypomethylated insertions frequently coincided with reduced proximal methylation. Representative examples included a hypermethylated intronic SVA in *B4GALT1* associated with increased surrounding methylation and a hypomethylated AluY near the *EFCAB13* promoter associated with reduced adjacent methylation (Fig. 2G and H). Together, these results indicate that MEIs both reflect and reshape local methylation landscapes, providing an epigenetic route through which insertions can influence nearby regulatory environments.

### TEs shape transcript architecture and generate polymorphic isoforms

We next asked whether poly-TEs contribute to transcriptome diversity by being incorporated into RNA isoforms. To define TE-containing transcript structures, we generated a high-confidence long-read transcriptome from 202 lymphoblastoid cell lines (LCLs), comprising 293,298 isoforms (fig. S3A to E). Approximately half of these isoforms contained at least one TE fragment (Fig. 3A and B). We classified TE-containing isoforms by the structural role of the embedded TE, including TE-derived transcription start sites (TSS), internal splice sites (ISS), transcript end sites (TES) and internal exonic elements (IEE). Events affecting transcript initiation, splicing or termination accounted for 66% of TE-containing isoforms, with ISS being the most prevalent class (Fig. 3B). TE-derived sequences generally accounted for only a small fraction of transcript length (Fig. 3C). Only 256 isoforms (247 from GENCODE and 9 from HPRC2) were initiated within a TE and had at least 80% of their sequence derived from that TE; 97% of these initiating elements belonged to evolutionarily old TE families. These findings indicate that TEs predominantly contribute modular sequence segments to host isoforms rather than generating autonomous TE-derived RNAs. TE-containing isoforms were enriched among novel isoforms absent from GENCODE, particularly intronic and antisense isoforms, as well as among lncRNAs and newly annotated genes (fig. S3F and G). The structural roles of TEs also varied by family: LTRs and SINEs were enriched for TE-derived TSSs, whereas SINEs were enriched in ISS and IEE classes (Fig. 3D and fig. S3H).

**Fig. 3.**
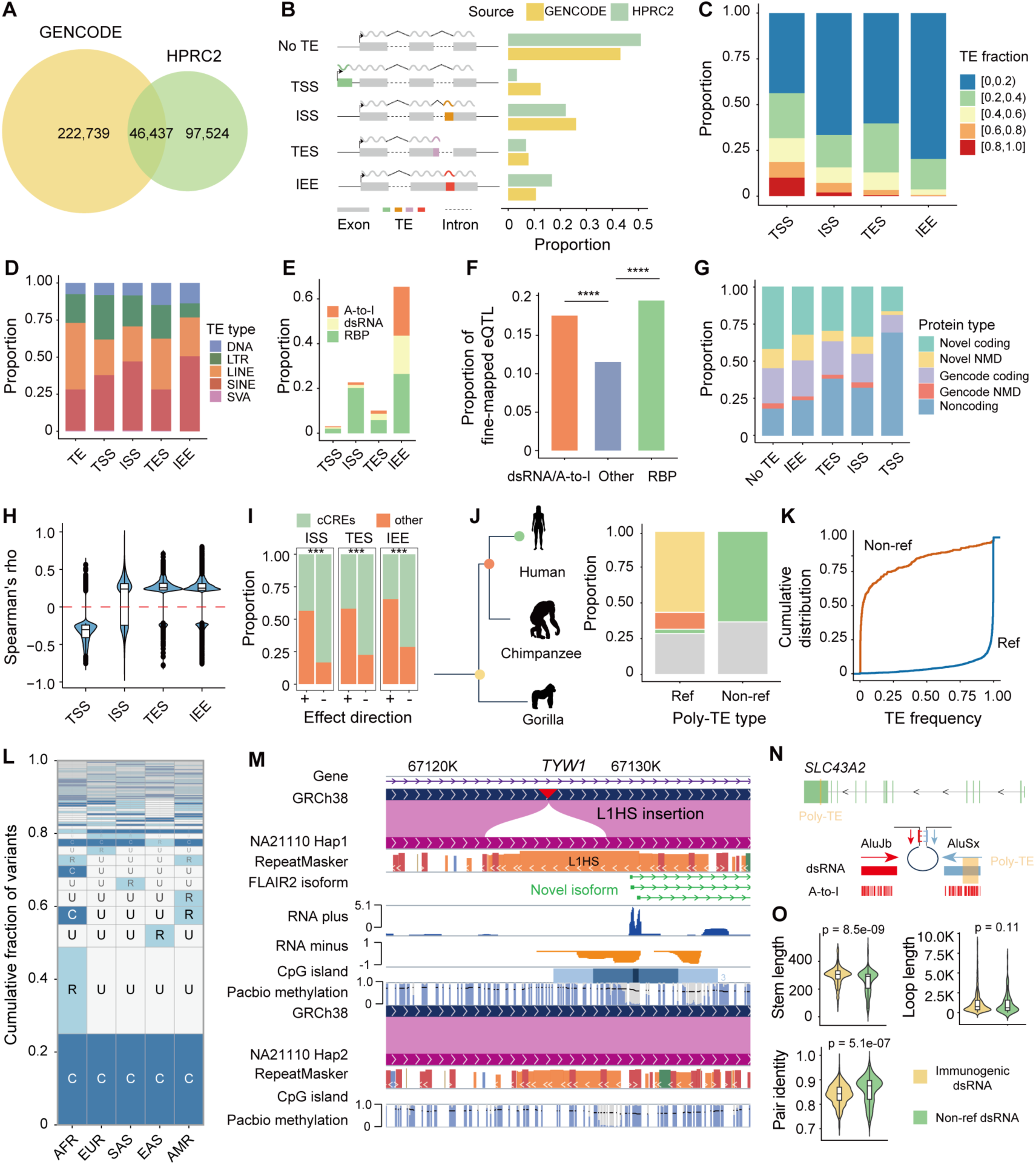
Polymorphic TEs remodel transcript architectures. (**A**) Number of TE-containing isoforms identified in HPRC2 and GENCODE (v47). (**B**) Proportion of isoforms assigned to TE-associated categories: no TE, isoforms without TEs; TSS, TE-derived transcription start site; ISS, TE-derived splice site; TES, TE-derived transcript end site; IEE, internal exonic elements, isoforms containing TE fragments that do not contribute to TSS, ISS or TES. (**C**) Fraction of isoform sequence derived from TEs. (**D**) Enrichment of TE types among isoform-associated TEs in HPRC2, compared with the reference-genome TE background. (**E**) Proportion of isoform-associated TEs assigned to RNA-regulatory categories. TEs overlapping multiple features were assigned hierarchically, with A-to-I RNA-editing sites prioritized over predicted double-stranded RNA (dsRNA) structures and RNA-binding protein (RBP) binding sites. (**F**) Proportion of internal exonic TEs with fine-mapped eQTLs from MAGE or GTEx (v10), stratified by overlapping with predicted dsRNA structures, A-to-I RNA-editing sites or RBP binding sites. P values were calculated by two-sided Fisher’s exact tests comparing internal exonic TEs with and without each feature. ****P < 0.0001. (**G**) Proportion of TE-containing isoforms with coding potential. NMD, nonsense-mediated decay. (**H**) Correlation between TE methylation and TE-containing isoform expression. TE-eQTMs were defined as associations between methylation levels of TEs and expression levels of their corresponding TE-containing isoforms. (**I**) Enrichment of negatively correlated TE-eQTMs in cis-regulatory elements (cCREs). Enrichment was assessed by two-sided Fisher’s exact tests comparing cCRE-overlapping TE-eQTMs with non-cCRE TE-eQTMs. ***P < 0.001. (**J**) Phylogenetic states of isoform-embedded polymorphic TEs (poly-TEs). Ref, reference poly-TEs; non-ref, non-reference poly-TEs. Species silhouettes were obtained from PhyloPic. Yellow, shared by human and gorilla; orange, shared by human and chimpanzee; green, human specific. (**K**) and (**L**) Allele-frequency and geographic distributions of isoform-embedded poly-TEs across 1000G superpopulations. (**M**) HPRC epigenome browser view of a L1HS insertion initiating a novel *TYW1* isoform, with tracks for gene annotation, sequence alignment, RepeatMasker, isoform structure identified by FLAIR2, Kinnex long-read RNA peak, CpG islands, and PacBio methylation. NA21110 Hap1 carries an intronic L1HS insertion (red arrowhead, boxed) absent from GRCh38 and NA21110 Hap2, which creates a hypomethylated CpG island and initiates three novel *TYW1* isoform. (**N**) Example of a poly-TE involved in a known immunogenic dsRNA. (**O**) Structural comparison between predicted dsRNAs formed by non-reference poly-TEs and known immunogenic dsRNAs.

TE incorporation also introduced RNA regulatory modules and altered the fate of host isoforms. Although IEEs do not by themselves define transcript boundaries, a subset located in UTRs, predominantly 3′UTRs, overlapped predicted double-stranded RNA (dsRNA) structures or RNA-binding protein (RBP) binding sites, and IEEs carrying these regulatory modules were significantly more likely to be fine-mapped eQTLs than other IEEs, suggesting that IEEs can exert regulatory effects through post-transcriptional mechanisms (fig. S3I and Fig. 3E and F). A subset also overlapped previously validated immunogenic dsRNAs(*38*), linking TE-derived IEE to RNA-duplex features with potential innate immune relevance. TE-containing isoforms were less often protein-coding than no-TE isoforms, especially those initiated from TE-derived TSS, whereas predicted nonsense-mediated decay (NMD) susceptibility was highest among IEE-containing isoforms. Nevertheless, a subset retained coding potential and contributed to predicted proteomic diversity (Fig. 3G).

TE methylation further provided an epigenetic layer linking incorporated TEs to isoform expression. Isoform level TE methylation–expression analysis identified 6,611 significant associations between TE methylation and isoform expression among 292,335 tested TE–isoform pairs. Methylation of TSS-associated TEs was predominantly negatively correlated with host isoform expression, consistent with promoter methylation-mediated repression, whereas gene-body TEs were more often positively correlated with isoform expression. Intriguingly, negatively correlated gene-body TEs were enriched for genic cCREs, indicating that TE methylation can influence isoform expression through both promoter-proximal and enhancer-like regulatory contexts (Fig. 3H and I).

We next focused on poly-TE-containing isoforms and asked whether polymorphic TEs generate transcript structures remained variable among present-day humans (table S3). We found 2,544 poly-TEs (112 MEIs and 2,432 non-MEIs) in the human reference GRCh38 incorporated into 11,183 isoforms (HPRC2 and GENCODE v47) and spanned all structural classes (fig. S3J). Among these, 452 poly-TEs were incorporated into 1,735 HPRC2 isoforms, and 2,394 poly-TEs were incorporated into 9,448 GENCODE v47 isoforms. Because most of these reference poly-TEs were shared with other apes and segregated at high allele frequency, their polymorphism largely reflects loss of established TE-derived transcript structures rather than recent TE gain (Fig. 3J and K). To capture non-reference poly-TE-containing isoforms, we further mapped long-read RNA-seq data to personal genome assemblies. This analysis identified 773 isoforms associated with 324 non-reference poly-TEs (175 MEIs and 149 non-MEIs). Most of these insertions were rare and human-specific (Fig. 3J and K), likely contributing to their lower representation among isoforms. Nevertheless, a substantial subset was common across superpopulations despite being absent from GRCh38 (Fig. 3L), highlighting transcriptomic variation missed by reference-based annotation.

Non-reference poly-TEs contributed to isoform diversity through multiple mechanisms. Among 31 non-reference TSS-providing poly-TEs, 10 showed haplotype-specific hypomethylation, suggesting that escape from epigenetic repression enables a subset of recent insertions to initiate transcription. All resulting isoforms were chimeric host–TE transcripts rather than autonomous TE-derived RNAs. Examples included an intronic AluY initiating an isoform of *HSD17B7*, additional TSS-associated insertions near *LYRM7*, *MYBL2*, and *ZFYVE19*, and an antisense full-length L1HS in *TYW1* associated with three novel isoforms, including two L1HS-initiated isoforms and one isoform incorporating L1HS sequence as the first exon (Fig. 3M and fig. S4). Consistent with this, RNA tracks revealed promoter-proximal divergent transcription around the L1HS insertion (Fig. 3M). Other non-reference poly-TEs introduced novel splice sites (*APOBEC3B*, *MUTYH*), contributed to transcript termination (*HSD17B12*, *CD109*) or appeared as IEEs within expressed isoforms (fig. S5), further illustrating the birth of TE-derived transcript structures that are invisible to the linear reference genome.

Finally, we examined whether poly-TEs could generate variable dsRNA-forming elements within expressed isoforms. In the reference genome GRCh38, 61 poly-TEs overlapped previously validated immunogenic dsRNAs(*38*) and 23 formed one arm of the paired dsRNA structure, indicating that deletion polymorphisms can remove or reshape known immunogenic dsRNA elements (Fig. 3N). Among non-reference poly-TEs incorporated into isoforms, 127 were predicted to form dsRNA structures with nearby pre-existing repeats in the same exon. Adenosine-to-inosine (A-to-I) editing within 32 insertions provides in vivo support for duplex formation, and their structural similarity to known immunogenic dsRNAs suggests that some may have functional consequences for RNA regulation or immune sensing (Fig. 3O). Together, these results show that TE sequences are widespread modular components of human transcript isoforms, and that poly-TEs reveal ongoing gain, loss and diversification of TE-derived transcript structures and RNA-duplex-forming elements. This variation provides a transcriptome-level basis for understanding how poly-TEs exert RNA level regulatory effects.

### Polymorphic TEs shape gene regulation through epigenetic tuning and isoform-specific RNA mechanisms

To systematically evaluate the molecular consequences of poly-TEs, we mapped cis-expression quantitative trait loci (eQTLs) in 201 HPRC2 LCL samples with graph-based genotypes and long-read transcriptome. Gene-level mapping identified 1,037 expression-associated TEs (eTEs) for 921 genes, whereas isoform-level mapping revealed a broader landscape, identifying 2,840 eTEs associated with over 8,000 isoforms (table S4). Fine-mapping with SuSiE(*39*) nominated 599 high-confidence candidate regulatory poly-TEs, only 20% of which were shared between gene- and isoform-level analyses, indicating that most effects were resolved only at isoform resolution. Complementary mapping in the larger MAGE LCL cohort (n = 731) recovered additional fine-mapped regulatory poly-TEs, including 301 fine-mapped eTEs and 143 splicing-associated TEs (sTEs) (Fig. 4A and fig. S6A, B).

**Fig. 4.**
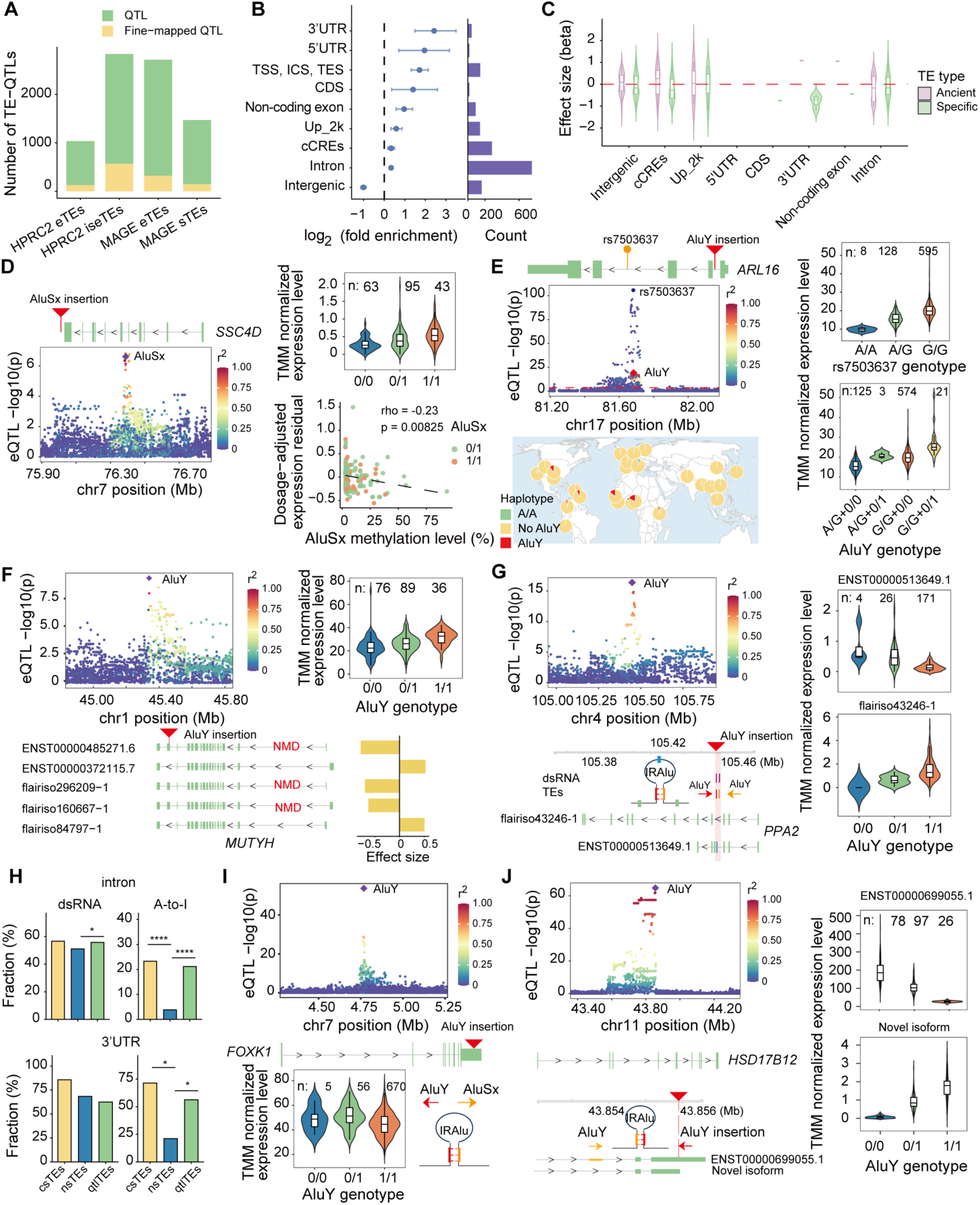
Molecular QTLs resolve the epigenetic and post-transcriptional mechanisms by which polymorphic TE alleles alter gene and isoform output. (**A**) Numbers of fine-mapped and non-fine-mapped gene-level TE-eQTLs (eTEs), isoform-level TE-eQTLs (iseTEs) and splicing TE-QTLs (sTEs) identified in the HPRC2 and MAGE datasets. eTEs, iseTEs and sTEs denote associations between polymorphic TE (poly-TEs) genotypes and gene expression, isoform expression and splicing, respectively. (**B**) Functional annotation enrichment of TE-QTLs. Enrichment was assessed by two-sided Fisher’s exact tests comparing TE-QTLs with distance- and frequency-matched non-QTL TEs. TSS, transcription start site; ISS, splice site; TES, transcript end site; Up_2k, 2-kb region upstream of genes. Error bars represent 95% confidence intervals. (**C**) Effect sizes of poly-TEs on gene expression across genomic and evolutionary contexts. Ancient poly-TEs are shared with outgroups, whereas human-specific poly-TEs are detected only in humans. (**D**) and (**E**) Fine-mapped eQTL signals for poly-TE insertions associated with *SSC4D* (D) and *ARL16* (E) expression. The map in panel (E) shows the geographic distributions of the two independent *ARL16* eQTL signals. (**F**) Fine-mapped QTL signals for AluY insertions associated with *MUTYH* isoform usage. NMD, nonsense-mediated decay. Isoforms were classified as NMD-prone under the 50-nt rule (premature stop codon > 50 nt upstream of the last exon–exon junction), using GENCODE biotypes for annotated transcripts and SQANTI3 predicted_NMD for novel ones. Effect size was from isoform-level eQTL. (**G**) Fine-mapped QTL signals for AluY insertions associated with *PPA2* isoform usage. IRAlu, inverted repeat Alus. (**H**) Enrichment of predicted dsRNA structures and adenosine-to-inosine (A-to-I) RNA-editing sites among intronic and 3’UTR TE-QTLs. Enrichment was assessed by two-sided Fisher’s exact tests comparing TE-QTLs with region-matched non-QTL TEs. csTEs, fine-mapped TE-QTLs; qtlTEs, TE-QTLs; nsTEs, non-QTL poly-TEs. (**I**) and (**J**) Fine-mapped eQTL signals for poly-TE insertions associated with *FOXK1* (I) and *HSD17B12* (J) expression. For (D) to (G) and (I) to (J), local Manhattan plots show QTL association signals, with focal poly-TEs highlighted as diamonds and point color denoting linkage disequilibrium (LD; r²) with the focal poly-TE; gene or isoform tracks show the poly-TE position; violin plots show TMM-normalized expression or isoform usage stratified by genotype. In (D), scatter plots show Spearman correlations between AluSx methylation and dosage-adjusted target-gene expression. In (E), fine-mapping identified two credible sets: rs7503637 and the AluY insertion.

These candidates were enriched in gene bodies and regulatory elements (Fig. 4B), with effect directions shaped by genomic context and evolutionary age (Fig. 4C and fig. S6C). Deletion of ancient TE-derived cCREs or exonic regions was frequently associated with reduced expression, consistent with long-term regulatory incorporation, whereas human-specific poly-TE insertions more often acted as disruptive alleles, particularly in exonic or cCRE contexts (Fig. 4C and fig. S6C). Thus, poly-TEs affect expression through both loss of ancient TE-derived elements and gain of new, often disruptive TE insertions.

At the DNA level, many regulatory effects were consistent with methylated TE alleles disrupting pre-existing regulatory architecture or remodeling nearby methylation landscapes. Human-specific TE-containing haplotypes within or near cCREs often showed reduced expression (Fig. 4C). Among loci with MEI-associated flanking methylation changes, 93 of them were associated with gene expression changes; most of them were hypermethylated (89 of 93), and 56 were associated with reduced target-gene expression. Representative loci included enhancer-region insertions at *B4GALT1* and *DGKE* and a promoter-proximal AluY insertion at *ZNF492*, all of which showed increased nearby methylation and reduced target-gene expression in carriers (figs. S6D and S7A and B). *ZNF492* and *DGKE* further showed methylation-sensitive repression among carriers, with higher methylation associated with stronger expression reduction (figs. S6D and S7A).

Conversely, we also found cases in which hypomethylation exposed TE-derived regulatory potential and was associated with increased gene expression. Among 28 consistently hypomethylated poly-TEs (hypomethylated in > 80% of all carriers) associated with differential gene expression, the presence of 24 of them were associated with increased gene expression, consistent with their regulatory potential as enhancers. The hypomethylated AluSx insertion downstream of *SSC4D* illustrated this methylation dependence: the insertion itself was associated with increased *SSC4D* expression, but among carriers, the *SSC4D* expression was negatively associated with AluSx methylation (Fig. 4D and fig. S8A). Poly-TE alleles can also add regulatory effects to pre-existing eQTL haplotypes. At *ARL16*, which encodes an innate immune regulator of RIG-I signaling, a hypomethylated African-specific AluY insertion occurred on a high-expression SNP haplotype and further increased transcription as an independent regulatory signal (Fig. 4E and fig. S8B). Together, these results indicate that the local methylation state determines the regulatory impact of TE insertions.

Isoform-resolved analysis further revealed isoform-level regulatory mechanisms that were obscured at the gene level. At *MUTYH*, an intronic AluY insertion moderately increased total gene expression, but isoform-level analysis showed that this increase reflected a coordinated shift away from NMD-prone isoforms toward non-NMD isoforms, together with de novo AluY-derived splice-site usage in a rare transcript (Fig. 4F and fig. S5). Thus, poly-TEs can alter the expression output not only by changing total gene expression, but also by redistributing expression across isoforms with distinct molecular fates. In other cases, poly-TEs directly became components of transcript architecture: an intronic insertion provided an alternative TSS and first exon for *TMEM68*, whereas an AluY insertion introduced a novel TES for *TRIM22* (fig. S8C and D).

One RNA-level mechanism involved intronic TE-derived dsRNA structures that remodel splicing(*40*). At *PPA2*, a sense AluY insertion in intron 4 is predicted to pair with a pre-existing antisense AluY in intron 3, forming an intronic dsRNA structure spanning exon 4. Consistent with the prediction, the insertion was associated with downregulation of exon 4–containing isoforms and a reciprocal increase in an exon 4-skipping isoform (Fig. 4G). Similar dsRNA configurations at *RSRC2* further support this as a recurrent mechanism for splicing diversification (fig. S8E). Genome-wide, intronic regulatory poly-TEs showed a modest increase in predicted dsRNA-forming potential and were significantly enriched for A-to-I RNA editing sites (Fig. 4H). Because A-to-I editing marks adenosine deaminase acting on RNA (ADAR)-engaged dsRNA in vivo, this enrichment further supports a broader contribution of poly-TE-derived RNA structures to splicing-associated regulatory variation.

Poly-TEs also acted through 3′UTR-associated RNA duplexes and post-transcriptional regulation. At *FOXK1*, oppositely oriented Alu elements within the 3′UTR were predicted to form an inverted-repeat Alus (IRAlu) duplex associated with reduced expression (Fig. 4I), consistent with established effects of IRAlu on RNA stability, localization and translation(*41, 42*). Notably, the polymorphic AluY also formed one arm of a previously validated immunogenic dsRNA(*38*). At *HSD17B12*, isoform-level resolution refined a previously reported association between a polymorphic 3′UTR AluY insertion and reduced expression(*16, 21*). Although the insertion increased usage of a novel isoform terminating within the AluY, the majority of the effect was the reduced usage of the canonical isoform (Fig. 4J and fig. S5), because the novel isoform represented only a small fraction of total expression. For the canonical isoform, the inserted AluY was predicted to pair with a pre-existing intronic AluY, suggesting that long-range 3′UTR–intron pairing may contribute to repression through RNA processing, stability or RBP recruitment(*43*). Genome-wide, 3′UTR eTEs were enriched for A-to-I RNA editing sites, supporting ADAR-engaged RNA structures as a broader feature of post-transcriptional poly-TE regulation (Fig. 4H). Together, these analyses show that poly-TEs reshape gene output through context-dependent effects on methylation, local cis-regulatory activity, transcript architecture and RNA processing.

### Regulatory polymorphic TEs provide molecular mechanisms underlying complex-trait associations

To link poly-TE-mediated regulation to broader phenotypic consequences, we asked whether these alleles underlie trait-associated haplotypes identified by SNP-based genome-wide association studies (GWAS). Although poly-TEs are rarely genotyped directly in GWAS cohorts, ∼80% of common poly-TEs were strongly tagged by nearby SNPs (r² > 0.8), especially MEIs and intermediate-frequency variants, whereas lower-frequency poly-TEs were poorly captured (Fig. 5A and fig. S9A), highlighting the value of our pangenome-resolved genotypes in identifying missing structural associations.

**Fig. 5.**
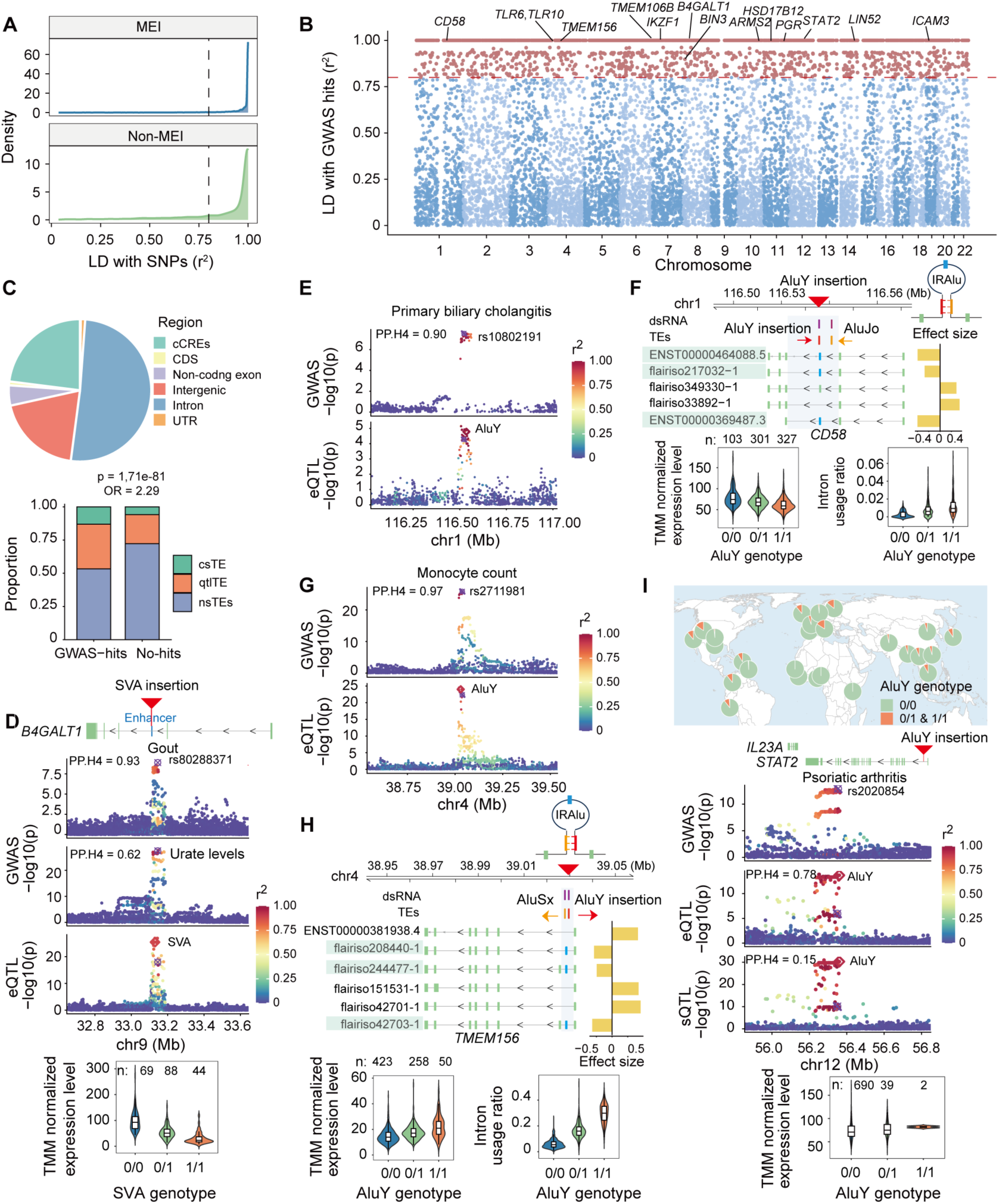
Polymorphic TEs resolve regulatory mechanisms at trait-associated loci. (**A**) Linkage disequilibrium (LD) between polymorphic TEs (poly-TEs) and nearby common SNPs (minor allele frequency > 5%). (**B**) Genomic distribution of poly-TEs linked to independent lead variants in the GWAS Catalog. (**C**) Genomic annotation and enrichment of trait-associated poly-TEs. csTEs, fine-mapped TE-QTLs; qtlTEs, TE-QTLs; nsTEs, non-QTL poly-TEs. Enrichment was assessed by two-sided Fisher’s exact tests comparing trait-associated and non-associated common poly-TEs. (**D**) Colocalization of urate-level and gout GWAS signals with a SVA insertion into an enhancer linked to *B4GALT1* expression. PP.H4, posterior probability of colocalization. (**E**) Colocalization of primary biliary cholangitis GWAS signal with an intronic AluY insertion linked to *CD58* expression. (**F**) The molecular mechanism underlying the eQTL and sQTL signal of *CD58*. IRAlu, inverted repeat Alus. Effect size was from isoform-level eQTL. (**G**) Colocalization of monocyte-count GWAS signal with an intronic AluY insertion linked to *TMEM156* expression. (**H**) The molecular mechanism underlying the eQTL and sQTL signal of *TMEM156*. (**I**) Geographic distribution of a Neanderthal-introgressed intronic AluY insertion associated with *STAT2* expression and colocalized with a psoriatic arthritis GWAS signal. For (**D**) to (**I**), molecular-QTL local Manhattan plots show variants associated with expression or splicing of the indicated target, with the focal poly-TE highlighted as a diamond, and point color denoting LD (r²) with the GWAS lead SNP. GWAS local Manhattan plots show variants associated with the indicated traits, with the lead SNP marked by a square-with-cross symbol. Gene or isoform tracks show the position of the focal poly-TE relative to the target gene or isoform. The shaded regions in (F) and (H) mark the focal introns. Violin plots show TMM-normalized gene expression or intron usage ratio stratified by poly-TE genotype.

Using GWAS Catalog lead variants(*44*) and ancestry-matched linkage-disequilibrium (LD) estimates, we identified 2,892 common poly-TEs in strong LD with GWAS signals, representing 16% of common poly-TEs (table S5 and Fig. 5B). These candidates were enriched in genic regions, cCREs and molecular QTLs compared with common poly-TEs not in strong LD with GWAS lead variants (Fig. 5C). Notably, 1,350 of these candidates were also eQTLs or sQTLs, and 368 were fine-mapped as putative causal TE-QTLs, establishing them as potential causal alleles for complex trait associations.

To resolve specific mechanisms, we prioritized loci where trait-linked poly-TEs colocalized with molecular QTLs. Our framework successfully recovered known associations while uncovering novel disease contexts and regulatory mechanism. For instance, at *B4GALT1*, a known enhancer-disrupting SVA previously implicated in autoimmune disease(*24*), also shape local epigenetic context and was linked here to urate level and gout GWAS signals, extending this regulatory TE locus to a metabolic disease context (Fig. 2G and Fig. 5D). Similarly, at *CD58*, an intronic AluY insertion has been long associated with multiple sclerosis risk and altered splicing(*19*), but how this insertion alters splicing has remained unclear. We found that this insertion was linked to primary biliary cholangitis and acted as a gene-level eQTL, isoform-level eQTL and sQTL for *CD58* (Fig. 5E and F). Mechanistically, the sense AluY insertion is predicted to pair with a pre-existing antisense AluJo to form an intronic dsRNA structure, providing a previously unrecognized mechanism for the exon 3 skipping and reduced expression observed at this risk locus (Fig. 5F).

Expanding beyond known loci, we identified previously unreported trait-associated regulatory candidates. An intronic AluY insertion in *TMEM156* was in strong LD (r^2^ = 0.92) with a monocyte-count GWAS signal and fine-mapped as both a gene-level and isoform-level eQTL and sQTL for *TMEM156*, linking the same allele to overall expression and isoform usage changes. Colocalization supported a shared association between the *TMEM156* eQTL and monocyte-count signal (Fig. 5G). Similar to the case of *CD58*, this insertion was predicted to form a dsRNA with a neighboring AluSx, providing a plausible explanation for altered splicing and downregulation (Fig. 5H).

Archaic introgression provided an additional source of trait-linked poly-TE alleles. At the *STAT2* locus, a Neanderthal-derived intronic AluY insertion linked to psoriatic arthritis was fine-mapped as both a *STAT2* eQTL and an *IL23A* sQTL, but only the *STAT2* eQTL colocalized with the disease association, implicating altered *STAT2* expression as the more likely mechanism (Fig. 5I). The AluY introduced an intermediately methylated CpG island and was associated with increased *STAT2* expression (fig. S9B). Across the genome, 25 archaic-derived poly-TEs were in strong LD with GWAS signals, indicating that introgressed TEs contribute a subset of present-day trait-associated haplotypes. Together, these findings show that poly-TE-derived regulatory alleles can shape human traits, providing sequence-resolved regulatory mechanisms for a subset of SNP-based association signals.

### Regulatory polymorphic TEs show signatures of recent positive selection in contemporary human populations

Given that poly-TEs generate widespread regulatory variation, we asked whether these alleles have been targets of recent positive selection. Using the linkage-based nSL (number of segregating sites by length) statistic(*45*), we identified 840 poly-TE loci with signatures of positive selection, including 615 MEIs and 225 non-MEI poly-TEs (table S6). These signals were highly structured across populations, with 406 population-specific and 693 superpopulation-specific events (Fig. 6A). Selection acted bidirectionally: 398 loci favored TE presence, whereas 432 favored TE absence, with 10 showing population-specific directionality. Both TE-presence- and TE-absence-favored alleles were younger than frequency-matched controls, particularly at high frequencies, consistent with positive selection accelerating their rise beyond neutral age–frequency expectations (Fig. 6B and fig. S10A). This bidirectionality suggests that the regulatory novelty introduced by TE can be either beneficial or deleterious depending on context. Consistent with this context dependence, an intronic L1HS insertion in *LEPR* was favored in two Chinese populations but disfavored in the Luhya population (LWK) (Fig. 6C).

**Fig. 6.**
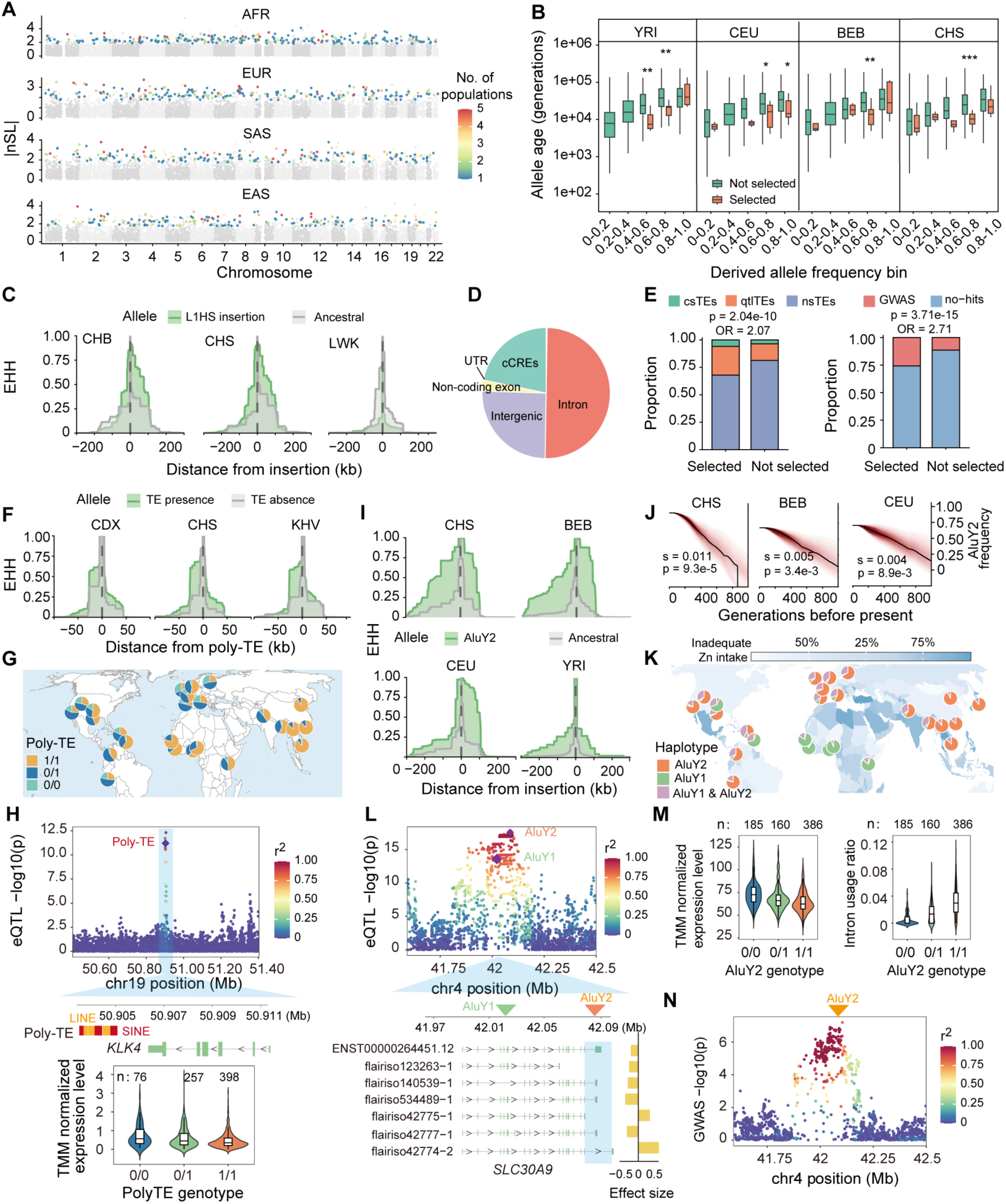
Recent selection on polymorphic TE-derived alleles across human populations. (**A**) Genome-wide distribution of absolute number of segregating sites by length (|nSL|) values for polymorphic TEs (poly-TEs), calculated in 20 populations and summarized by 1000G superpopulation. Gray, not selected; colored, putatively adaptive, with color scale indicating the number of selected populations. For poly-TEs selected in multiple populations within a superpopulation, the highest |nSL| is shown. (**B**) Derived allele ages of TE-presence-selected poly-TEs and frequency-matched non-selected controls. Ages are plotted on a log_10_ scale. Within each population and frequency bin, selected and non-selected poly-TEs were compared using two-sided Wilcoxon rank-sum tests; asterisks denote significance: *P < 0.05, **P < 0.01 and ***P < 0.001. (**C**) Extended haplotype homozygosity (EHH) around an intronic L1HS insertion in *LEPR*. Population-specific EHH patterns indicate recent haplotype expansion around this locus. Population abbreviations follow 1000G nomenclature. (**D**) Genomic annotation of putatively adaptive poly-TEs favoring TE presence. (**E**) Enrichment of adaptive poly-TEs favoring TE presence among molecular QTLs and GWAS-linked variants. csTEs, fine-mapped TE-QTLs; qtlTEs, TE-QTLs; nsTEs, non-QTL poly-TEs; no-hit, non-GWAS-linked poly-TEs. Enrichment was assessed by two-sided Fisher’s exact tests comparing adaptive and non-adaptive poly-TEs. (**F**) EHH around an ancestral poly-TE downstream of *KLK4*. (**G**) and (**H**) Geographic distribution and eQTL evidence linking the poly-TE to *KLK4* expression. Genotypes are coded as TE presence (1) or absence (0). (**I**) EHH around AluY2, a potentially adaptive AluY insertion in the last intron of *SLC30A9*. (**J**) Inferred derived-allele frequency trajectories of AluY2; s, selection coefficient; P, likelihood-ratio test P value; shading, posterior intervals. (**K**) Geographic distribution of the AluY2 allele shown alongside country-level prevalence estimates of inadequate zinc intake from the 2018 national nutrient intake inadequacy dataset (https://raw.githubusercontent.com/cfree14/global_intake_inadequacies/main/output/2018_national_nutrient_intake_inadequacy_estimates.csv). (**L**) QTL evidence linking AluY2 to *SLC30A9* expression and isoform usage. Effect size was from isoform-level eQTL. (**M**) TMM-normalized *SLC30A9* expression and intron usage ratio stratified by AluY2 genotype. The focal intron was marked in (L) as shaded region. (**N**) LD between AluY2 and depression-associated variants near *SLC30A9*.

The regulatory relevance of these selected alleles is underscored by their genomic context: among the 398 selection-favored TE-presence alleles, 74.4% resided within genic regions or cCREs (Fig. 6D). Notably, 127 were eQTLs or sQTLs, including 22 fine-mapped eTEs or sTEs, and selected poly-TEs were significantly enriched for QTLs and GWAS-linked variants compared to non-selected counterparts (Fig. 6E), suggesting them as likely causal drivers rather than passive markers of adaptive haplotypes. This adaptive repertoire also includes archaic-derived variation, such as four selected AluY insertions introgressed from archaic hominins, including an intronic AluY in *AUH* favored in East Asian populations (fig. S10B).

To resolve specific adaptive mechanisms, we examined a 1,576-bp composite SINE/LINE poly-TE located 1,276 bp downstream of *KLK4*, a serine protease required for enamel maturation and implicated in epithelial biology. An expression-reduced *KLK4* haplotype was previously shown to have undergone adaptive selection in East Asians(*46*), but the underlying mechanism remained unresolved. Our results support the poly-TE as a potential functional driver. The poly-TE near *KLK4* showed strong adaptive selection signals in CDX, CHS and KHV, matching its high frequency in East Asia (Fig. 6F and G). Because the TE sequence was shared with outgroups, this signal likely reflects selection maintaining or increasing an ancestral TE-containing haplotype rather than selection on a new insertion. Fine-mapping identified this variant as a functional eTE, with TE presence associated with decreased *KLK4* expression (Fig. 6H). Mechanistically, the poly-TE introduced a heavily methylated CpG island, whereas the TE-absent haplotype lacked this CpG island and showed low methylation (fig. S10C), suggesting that the poly-TE may represent a methylated and repressive regulatory module that were lost in some human populations.

Whereas *KLK4* illustrates selection on an ancestral TE-containing haplotype that modulates expression through methylation, *SLC30A9* provides a contrasting example in which selection favored an insertion-bearing haplotype affecting transcript processing. The zinc transporter gene *SLC30A9* has been implicated in adaptation to micronutrient availability(*47, 48*), but the underlying mechanism has been elusive. Two intronic AluY insertions in *SLC30A9* showed strong adaptive selection signals across non-African populations, but sweep signals consistently favored the presence of AluY2, located 59 bp upstream of the terminal exon, and the absence of AluY1 in intron 4 (fig. S10D and Fig. 6I and J). The two insertions define a population-dependent haplotype background that is largely separated in African and East Asian populations but more mixed in Europeans, so the sweep signal alone cannot distinguish selection for AluY2 from selection against AluY1 (Fig. 6K).

Several lines of molecular evidence support AluY2 as the more likely functional driver. Although both insertions were associated with *SLC30A9* expression and splicing, AluY2 was included in the sQTL credible set and is positioned immediately adjacent to the terminal splice region (Fig. 6L and M). Conditional genotype analysis further separated their effects on transcript processing: within fixed AluY2 genotype backgrounds, AluY1 had little effect on intron usage near the terminal exon, whereas AluY2 was consistently associated with terminal intron usage within matched AluY1 genotype backgrounds (fig. S10E). Consistent with this local effect, AluY2 promotes the usage of non-canonical isoforms at the expense of the canonical transcript (Fig. 6L and M), suggesting a splicing-mediated expression decrease. AluY2 was also in strong LD (r^2^ = 0.83) with a depression GWAS signal, linking cellular zinc homeostasis to complex behavioral traits (Fig. 6N). Together, these cases show that positive selection can act on both ancestral TE-containing haplotypes and derived insertion-bearing haplotypes, highlighting poly-TE-mediated tuning of methylation, expression and transcript processing as a substrate for ongoing human regulatory evolution.

## Discussion

Our study provides a sequence-resolved view of how TE-derived alleles that segregate among present-day humans can contribute to contemporary regulatory evolution. This layer of variation has been difficult to resolve because linear reference genomes poorly represent non-reference TE insertions, while short-read sequencing rarely capture the full sequence architecture of TE-containing structural variants(*14, 33*). Consistent with this limitation, nearly half of the HPRC2-resolved MEIs were absent from short-read call set. Beyond detection itself, short reads provide limited access to the inserted sequence, obscuring TE-derived variants that lack canonical signatures of recent mobilization(*30, 32, 49*). Resolving these alleles in haplotype-resolved pangenomes therefore changes the picture qualitatively, not just quantitatively. Human poly-TEs are not a single class of recent mobilizations, but a heterogeneous spectrum of TE-derived haplotypes arising through recent mobilization, structural remodeling of older repeats, ancestral segregation and archaic introgression, and subsequently shaped by demography and selection. Pangenome resolution thus does more than expand the catalog of TE variation; it recasts present-day poly-TE diversity as both a composite record of human evolutionary history and a sequence-rich substrate for ongoing regulatory evolution.

Our data further show that the regulatory potential of a polymorphic TE is conditional rather than intrinsic. At the DNA level, TE methylation both reflected local chromatin context and reshaped nearby methylation landscapes, while consistently hypomethylated insertions were enriched in promoters and cCREs. Thus, methylation helps determine whether a TE allele remains silent, perturbs nearby regulation or exposes cis-regulatory potential. Long-read transcriptomics extended this view to RNA output, identifying poly-TE-containing isoforms absent from linear references and revealing haplotype-specific transcript structures(*50*). These data show that TE-derived regulation extends beyond simple promoter activity or exonization(*9, 10, 51*): internal exonic TEs represent an important mode of TE incorporation into isoforms and can carry embedded regulatory modules. Furthermore, poly-TE-derived dsRNAs suggest that inter-individual variation in RNA immunogenicity can arise from the presence or absence of the dsRNA-forming element itself, providing a molecular basis for autoimmune disease risk(*38*).

Building on this epigenetic and transcriptomic map, molecular-QTL analyses connected latent TE-derived regulatory potential to measured, allele-specific effects on gene output. These effects were often not simple consequences of insertion presence alone. At the epigenetic level, loci such as *DGKE* further indicate that methylation variation among carriers can modulate the magnitude of expression effects, suggesting that the same TE allele can have different regulatory outputs depending on its chromatin context. At the gene level, many poly-TE associations were isoform-specific and directionally heterogeneous, indicating that gene-level eQTLs can average away effects on transcript initiation, splicing, termination or isoform fate(*52, 53*). At the RNA level, dsRNA-associated splicing remodeling and 3′UTR inverted-repeat effects suggest that polymorphic TEs can act through post-transcriptional mechanisms that depend on pairing partners and transcript contexts created by the allele. These findings extend prior examples of dsRNA-mediated regulatory change, including fixed innovations such as *TBXT*(*54–56*) and polymorphic SVAs at *ASIP*(*29*), to a recurrent, trait-relevant class of segregating TE alleles with molecular-QTL support. Thus, poly-TE effects can be hidden from conventional analyses not only because the underlying alleles are structurally difficult to resolve, but also because their consequences depend on isoform, RNA-structural and epigenetic context. Resolving these alleles on assembled haplotypes makes this conditional layer of regulatory variation mechanistically interpretable.

These findings further argue that polymorphic TEs represent a poorly captured component of complex-trait genetics. Because most GWAS do not directly test poly-TEs(*16, 23, 24*), their effects are often projected onto linked SNPs, obscuring the molecular allele responsible for the association. Our integration of pangenome-resolved poly-TEs with molecular QTLs and GWAS loci shows that such variants can provide sequence-resolved regulatory mechanisms for trait-associated signals, including effects on expression, splicing, methylation and RNA structure. Extending association studies to directly genotype or impute and test these structurally complex alleles should therefore improve mechanistic interpretation of GWAS and may recover a fraction of missing heritability that has remained inaccessible to SNP-centric analyses(*57*).

Placed in an evolutionary framework, these regulatory effects show that poly-TE variation is not merely a molecular phenomenon but part of standing regulatory variation exposed to constraint and selection. Although most poly-TEs are neutral or deleterious, and TE exaptation is usually recognized only after ancient insertions have fixed and bear cross-species conservation, our analyses identify a subset of still-segregating poly-TEs with signatures of recent, often population-specific, positive selection, molecular QTL effects and trait-associated haplotype links. Selection was bidirectional, favoring TE-present alleles in some populations and TE-absent alleles in others, as illustrated by population-dependent favored states at *LEPR*. Moreover, adaptive TE-associated variants need not arise only from newly derived insertions: the *KLK4* locus shows selection on an ancestral TE-containing haplotype that remains polymorphic. Thus, poly-TE-associated regulatory evolution can proceed through both new TE-derived activity and selection on pre-existing TE-containing haplotypes, linking innovation, constraint and local adaptation within the same evolutionary process.

Our study provides a framework for treating poly-TEs as sequence-resolved regulatory alleles whose evolutionary origins, molecular consequences and population trajectories can be interpreted together. Extending this framework to larger and more globally representative pangenomes(*32, 58*), biobank-scale molecular cohorts(*59*), diverse tissues(*60*), developmental stages and environmental perturbations will define the breadth and context specificity of poly-TE effects, while targeted functional validation will help distinguish causal mechanisms from linked haplotypic associations. Because the effects of poly-TEs are context dependent, future studies should jointly resolve TE sequence, regulatory context and phenotype. Together, our work shifts the study of TE-derived innovation from a retrospective account of ancient exaptation toward a population-level view of regulatory innovation still unfolding in present-day humans.

## Acknowledgements

We would like to thank Jana Elber for generously providing the PanGenie-genotyped 1000G samples generated using the HPRC2 pangenome graph.

## Author Contributions

T.W., J.J. and X.Y.Z. conceptualized the study. J.J., X.Y.Z., R.H.L., Z.D. and W.J.Z. performed the study. J.J. and X.Y.Z. wrote the manuscript. All authors read and revised the manuscript. T.W. supervised the study.

## Data, code and materials availability

All primary HPRC2 datasets used in this study, including haplotype-resolved genome assemblies, graph-derived variant calls, CpG island annotation, DNA methylation, long-read RNA-seq and TE annotations are publicly available through the HPRC data portal (https://data.humanpangenome.org).

High-coverage 1000 Genomes Project (1000G) short-read structural variant calls were obtained from (https://www.internationalgenome.org/data-portal/data-collection/1000genomes_30x). Poly-TE genotypes for 1000G samples were obtained from the HPRC2 PanGenie genotype release on Zenodo (https://zenodo.org/records/16737952). The GRCh38 reference genome was used (https://www.ncbi.nlm.nih.gov/datasets/genome/GCA_000001405.15/), gene and transcript annotations were obtained from GENCODE (v47) (https://www.gencodegenes.org/human/release_47.html). The hg38–panTro6 and hg38–gorGor6 reciprocal-best chain files were downloaded from the UCSC Genome Browser. MAGE RNA-seq data, gene expression matrices, splicing matrices, sample metadata, covariates and cis-eQTL/sQTL summary statistics were obtained from the MAGE public data release (accession: PRJNA851328, https://zenodo.org/records/10535719). GTEx QTL summary statistics (v10) were downloaded from the GTEx Portal (https://www.gtexportal.org/home/downloads/adult-gtex/qtl). Cis-regulatory element annotations were obtained from ENCODE SCREEN cCRE resources (https://screen.wenglab.org). RBP-binding site annotations used in this study are publicly available from RBP-Tar, an ENCODE eCLIP-derived database of experimentally supported RBP-binding sites (https://rbp-tar.biodata.ceitec.cz/download_all). A-to-I RNA editing sites were obtained from REDIportal V3.0 (https://rediportal.cloud.ba.infn.it/atlas/index.html). Predicted dsRNA in GRCh38 were obtained from dsRNAscan (https://dsrna.chpc.utah.edu). GWAS summary statistics and harmonized GWAS summary statistics were obtained from the GWAS Catalog (https://www.ebi.ac.uk/gwas).

Processed datasets generated in this study are available at (https://github.com/twlab/HPRC2_TE). Source data are provided with this paper.

Code used for the analyses presented in this paper is available on GitHub (https://github.com/twlab/HPRC2_TE)

## Supplementary Materials

Materials and Methods

Figs. S1 to S10

Tables S1 to S6

References (*61–96*)

